# The development of social interactions in *Corydoras aeneus* larvae

**DOI:** 10.1101/455188

**Authors:** RJ Riley, T Roe, ER Gillie, NJ Boogert, A Manica

**Affiliations:** Department of Zoology, University of Cambridge, CB2 3EJ; Cardiff University, The Sir Martin Evans Building, Museum Ave, Cardiff CF103AX; University of Exeter, Stocker Road, Exeter EX4 4QD

## Abstract

Very young animals develop life skills as they mature, and for social animals this includes the acquisition of social abilities such as communication. Many animals exhibit changeable patterns of social behavior based on development, and social experience during the juvenile period can be vital for the development of necessary social behaviors in adulthood. We investigated the development of a distinctive tactile interaction behavior in *Corydoras aeneus*, the Bronze Cory catfish. Adults use this behavior to coordinate group activities during foraging and flight responses from predators, and the development of this behavior in larvae is of interest in investigating how communication and social behaviors develop as an individual matures, and which factors affect their development. We found that larvae respond to applied tactile stimulation with a flight response far less often as larvae matured, implying that larvae become less sensitive to tactile stimulation with age. Given that adults frequently interact with one another tactilely, this development is consistent with developing appropriate social behavior in adulthood. We also found that social exposure affects the development of the larval response to tactile interactions with conspecifics, and that isolation in the earliest larval stage leads to a greater likelihood of responding to a tactile interaction with a conspecific with a flight response. This suggests that social exposure is important for developing an appropriate response to tactile stimulation in social settings and underscores the particular importance of early life experiences in the development of sociality.

## Introduction

Group living confers a number of advantages to social animals, including a reduced risk of predation (Magurran, 1990) and increased foraging efficiency (Pitcher, 1982). In order for social animals to reap the benefits of group living, individuals must coordinate their behavior to their group-mates’. Many social animals, however, begin their lives as undeveloped juveniles whose behavior gradually adapts to social interactions over time. For example, the labyrinth fish develops the ability to communicate vocally later in development than the ability to perceive conspecific vocalizations (Wysocki, 2001), and birds develop the ability to sing first in a generalized learning stage and later in development in a more specific learning stage (Irwin, 1988). The progression of social skills as individuals grow and mature has important consequences for group coordination.

Many types of group coordination, including cohesion, develop in groups alongside physical development in individuals. In the catfish *Corydoras paleatus*, the development of group cohesion and aggregative behaviors is highly correlated to anatomical developmental stage (Rodríguez-Ithurralde, 2014). The development of cohesive group behavior with age has also been documented in zebra fish, which begin to form cohesive shoals as individuals age (Buske and Gerlai, 2011). In Florida scrub jays, mobbing behavior, a group behavior used to deter predators, first appears in the fledgling stage and gradually develops into adult mobbing behavior over about three months (Francis, 1989).

The development of behavior over a young animal’s maturation to adulthood is vital for individuals and groups, and the social environment an animal experiences during development can have a profound impact on the social behaviors it exhibits as an adult (Slagsvold, 2002). This is apparent in species that experience imprinting, as when very early experiences determine mate preferences in great tits (Slagsvold, 2002). In zebrafish, a preference to shoal with conspecifics with particular coloration patterns is highly influenced by the patterns of the group-mates that an individual was raised with (Spence and Smith, 2007). The way social behavior can be determined by early experiences highlights how the mechanisms of development can have cascade effects on an animal’s behavior through its life. In rats, for example, individuals who experience early social isolation subsequently develop a behavioral condition which shares core features with schizophrenia (Fone and Porkess, 2008). Furthermore, many species have ‘critical periods’ in which developing individuals must experience specific stimuli in order to develop adult behaviors. Examples include language development in humans, in which individuals must be exposed to language within a window of development to fully develop language abilities (Kuhl, 2005) and song tuition in zebrafinches, which must occur within a specific period of development for a young bird to properly learn to sing (George, 1995).

Due to the importance of social development in group-living animals, the development of communication between individuals is of particular interest, as communication and interactions between individuals is a driving factor of group coordination in many species (Conradt and Roper, 2005). Adults of our study species *Corydoras aeneus*, the Bronze Cory catfish, utilize tactile interactions that seem to facilitate group coordination (Riley, 2018a) and mediate flight responses from a potential threat (Riley, 2018b). The usage of tactile interactions in the Bronze Cory catfish is unusual, as tactile stimulation often triggers a stereotyped threat response in fish called a ‘c-start’ that is deployed involuntarily when an individual perceives an urgent potential threat (Kimmel, 1980). The c-start threat response was first reported in detail by (Weihs, 1973) in trout and pike and is characterized by a two stage motor pattern (Figure 1). Stage 1 lasts 15–40 msec and is characterized by the ipsilateral contraction of axial muscle on one side of the body (Eaton and Emberley, 1991). The fish orientates away from the threat and resembles a ‘C shape’ from above at the end of stage one. The head and caudal fin flap lie in the same direction. Stage 2 is characterized by the straightening of the axial skeleton and acceleration caused by the lateral side of the fish displacing water (Eaton, 1988). This allows the fish to be propelled forwards in an escape trajectory. The underlying neural command appears to be ballistic, and the reflex is unaffected by sensory information once it is initiated (Eaton and Emberley, 1991). The fact that the Bronze Cory catfish interact tactilely with conspecifics without a c-start response suggests that Brone Cory catfish must either have an innate trait that stops them from interpreting tactile stimulation as a potential threat, or that they must develop a tolerance to tactile stimulation during development in order to interact with conspecifics.

**Figure 1.**
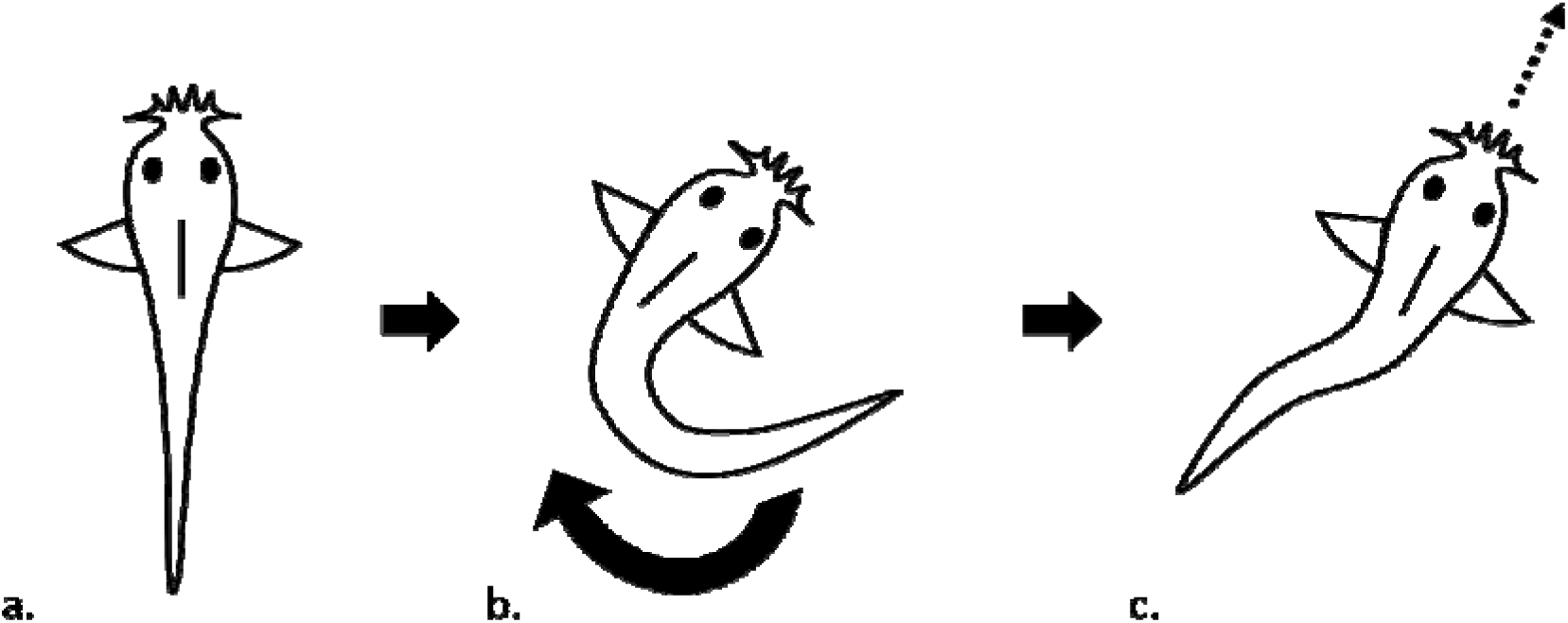
C-Start Schematic Diagram – Dorsal View. a.) Larva pre C start b.) Stage 1 – larva adopts ‘C shape’ during the fast phase. c.) Stage 2 – larva straightens body and is propelled forwards in an escape trajectory

Furthermore, the Bronze Cory catfish provides a unique study system for investigating the development of social interaction, as patterns of tactile interactions can be observed and responses to tactile interactions can be clearly recorded. This allows us to ask questions about the way individual larvae respond to tactile interactions during development, and about the role of social exposure in developing a tolerance to tactile stimulation. How do individuals develop the ability to interact and coordinate with others, and how do their experiences with conspecifics during early development affect their responses to tactile interactions later on?

We conducted two experiments to investigate the development of communication and sociality in larval Bronze Cory catfish. First, we investigated the ontogeny of the response to tactile stimuli by stimulating individuals within stable groups over regular intervals in the developmental period. We predicted that older larvae would respond to the tactile stimulation with a c-start less often than younger larvae. We also predicted that individuals that did respond to the tactile stimuli would be more likely to initiate tactile interactions later in development. Second, we investigated the effect of isolation on sociality and communication in larvae that ranged from 14 to 21 days post hatching (dph) in age. We predicted that isolated larvae would spend less time together, participate in fewer tactile communication interactions and be more likely to respond to tactile interactions with conspecifics with a c-start.

## Methods

### Study subjects

We obtained *C. aeneus* from three local pet shops in Cambridgeshire: Maidenhead Aquatics Cambridge, Pet Paks LTD, and Ely Aquatics and Reptiles. Fish in both experiments were maintained in reverse osmosis (RO) water purified to 15 or less total dissolved solids (TDS) and re-mineralized to 105-110ppm TDS using a commercially prepared RO re-mineralizing mix (Tropic Marin Re-mineral Tropic). The fish experienced a 12:12 light:dark cycle under fluorescent ambient room lighting at a temperature of 23 ± 1 °C. We can be certain that all eggs are from the same clutch are full siblings due to the unique sperm drinking behavior of females (Kohda, 1995).

### Development of Response to Tactile Stimulation

This experiment was done in two batches, with two weeks in between hatching. Eggs from three clutches per batch were hatched, and larvae were placed with full siblings in groups of five into a transparent, 21L plastic fish tank at 12 days post hatching (dph). Each tank had a small air driven aquarium filter for biological filtration. Larvae were fed “Interpet Liquifry Number 1” and “Interpet liquifry Number 3” twice daily. Aufwuchs from adult Bronze Cory tanks was introduced for larvae to graze on. In addition, larvae were fed twice a week with live *Panagrellus redivivus*. Larvae were always fed after experimentation on days when they underwent tactile stimulation. Larvae were stimulated tactility on 14, 24, and 31 dph, and their responses were filmed from a dorsal perspective (i.e. from above). The order in which the groups were filmed being stimulated was randomly determined. To optimize filming, before tactile stimulation commenced, the water level in the tanks was reduced to 11cm, and the air driven filter removed as was any detritus on the base of the aquarium. The larvae were left for 15 minutes to acclimatize.

A 25cm long glass rod which tapered to a 1mm blunt end was used to tactilely stimulate the larvae. Tactile stimulation was applied by RJR and TR, and stimulation was performed in rounds. Each of the five fish in a group was stimulated in turn during a round before the next round commenced. Rounds were separated by a period of at least 60s to avoid stressing the larvae excessively. Stimulation was standardized (Figure 3). A test cohort was used to develop the tactile stimulation protocol, from which no data were collected, before the experiment began. This ensured that the tactile stimulations were consistent and reproducible when data were collected. Each group of larvae underwent 10 or 11 rounds of tactile stimulation with each individual receiving tactile stimulation at maximum once per round. Stock tanks containing full siblings for all the experimental groups were maintained under identical conditions so that any deceased larvae could be replaced with full siblings. Over the course of the experiment, 11 larvae died (of either apparent deformities, or no obvious cause) in seven different groups, and were replaced with full siblings from the same clutch at least 24 hours before tactile stimulation was applied.

We scored the immediate reaction of a fish to tactile stimulation with a glass rod and recorded whether a tactile stimulation event led to that fish initiating a tactile interaction with a group-mate(s). We defined the immediate reaction as occurring within the first second after stimulation. Fish either “ignored”, c-started or performed a non-c-start flight response. A c-start was scored if a clear c-shaped body morphology was observed in conjunction with rapid movement. A response was scored as “ignored’ if the larva being stimulated remained static. A non-c-start flight response was scored if the larva did not remain static in response to tactile stimulation but also did not undergo a c-start. If a larva responded with a flight response of any sort (c-start or non-c-start), we also recorded if it subsequently initiated a social interaction with another larva within 5 seconds of the initial tactile stimulation.

An ethogram was developed to score the tactile stimulation of larvae filmed dorsally. Videos where scored by two researchers. Both researchers observed one full session of tactile stimulation (10 rounds of stimulation) together to develop scoring protocol for an ethogram. Four training videos were then scored independently by both researchers. Scores were compared, and inconsistencies resolved by referring to the scoring protocol and reaching a consensus about each interaction. A further 2 videos were scored independently and the scores of each measure were compared. All measures (immediate reaction, initiates an interaction, incidental or non-incidental) were within 90% agreement between both scorers and so the scoring was assessed as being consistent. The remaining videos were randomly divided up between the two researchers and were scored independently.

### Effect of Isolation on Sociality and Communication

To investigate the effect of social isolation on sociality and communication, larvae were reared in groups of 3-5 or in isolation until testing. All larvae were raised in identical mesh enclosures of roughly 9.5cm diameter suspended in 20L aquariums equipped with an Interpet MINI filters. Each 20L aquarium held 2-6 mesh enclosures. Larvae were fed twice daily with “Interpet Liquifry Number 1” and “Interpet liquifry Number 3” daily. Aufwuchs from adult Bronze Cory tanks was introduced for larvae to graze on. Larvae were tested when they reached the age range of 14-21 dph. and tested with the age range of 14-21 dph. After a 30 minute acclimatization period, larval behavior was filmed from dorsally for a 1-hour period. The camera was fixed 30cm above the water surface. There was no pseudo replication between videos.

We defined a tactile interaction as an event when one fish physically touches another. Interactions were only scored if both fish were on the bottom of the enclosure. Larvae were scored as together if they were within 2 body lengths of each other using the body length of the smallest fish in that group. Time was recorded as soon as two fish were within this distance and were on the bottom. Number of fish in the group was also recorded. Togetherness (ie cohesion) was terminated if one fish began side fixating having been on the bottom of the enclosure prior to that. If a fish briefly moved more than two body lengths away from its partner(s) but returned to within 2 body lengths in less than three seconds, then this was scored as one continuous period of togetherness without a break for the brief separation period. Periods of togetherness were only scored if individuals were together for at least 3 seconds.

Each tactile interaction had an initiator and a receiver. An initiator is the individual whose movement resulted in the interaction. A receiver is the individual who was touched by the movement of the initiator. We recorded the response of initiators and receivers to tactile interactions. A c-start was scored if a clear c-shaped body morphology was observed in conjunction with rapid movement, as is characteristic of the fast phase of the c-start response.

To assess the consistency of scoring protocol, we developed an ethogram to score the interactions between group-members. Videos where scored by RJR and TR. Scoring always began 30 minutes after the start of filming. Both researchers observed an hour of randomly selected footage together to develop scoring protocol for an ethogram. Three training videos were then scored independently by both researchers. Scores were compared, and inconsistencies resolved by referring to the scoring protocol and reaching a consensus about each interaction. A further three videos were scored independently and the scores of each measure were compared. All measures (total time together, total number of interactions, total initiator c-starts, total receiver c-starts, total spontaneous c-starts) were within 90% agreement between both scorers and so the scoring was assessed as being consistent. The remaining videos were randomly divided up between the two researchers and were scored independently. One researcher assigned codes for the videos so that the other could score them blind.

### Data analyses

We used a General Linear Model (GLM) with quasibinomial error structure to assess the differences between larvae ages 14, 24, and 31 dph in the proportion of tactile stimulation events that resulted in a c-start and that resulted in fish not responding (or ‘ignoring’) the stimulus. Similarly, we used a GLM with quasibinomial error structure to assess the proportion of tactile stimulation events where an individual responded to the stimulus in which the response involved a tactile interaction with another individual. For all analyses, we used group ID as a blocking factor.

We used a GLM with a Poisson error structure to assess the differences between socially-housed and isolation-housed groups in the total interactions each group underwent. We used a linear model (LM) to assess whether socially-housed and isolation-housed groups differed in the amount of time individuals were in proximity to one another. For all analyses, we used group ID as a blocking factor.

Statistical analysis was performed in R version 3.4.2.

## Results

### Development of response to tactile stimulation

Bronze Cory catfish larvae underwent a behavioral transformation in response to applied tactile stimulation starting from hatching and continuing until about 31 dph. Larvae responded to tactile stimulation with a c-start response at lower rates as they mature (quasibinomial GLM: F_1,19_= 61.8, p<0.001, Figure 1a). Furthermore, as larvae matured, they were significantly more likely to respond to tactile stimulation by remaining stationary and ‘ignoring’ the stimulus (quasibinomial GLM: F_1,19_= 44.5, p<0.001, figure 1b)

**Figure 1:**
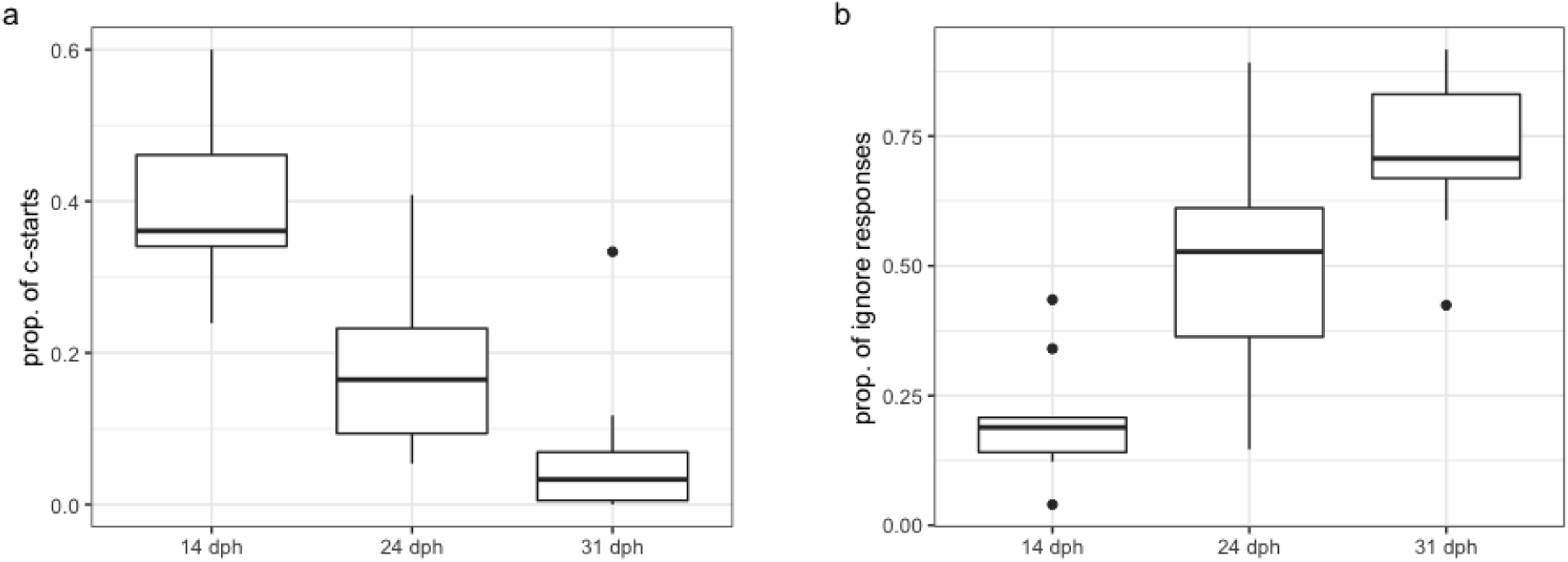
a) the proportion of tactile stimulation events that resulted in a c-start response in fry days 14, 24, and 31 dph; b) the proportion of tactile stimulation events that results in an ignoring response on days 14, 24, and 31 dph.

In addition, fry that do respond to a tactile stimulation event were much more likely to nudge their group-mates during their response. As fry mature, the proportion of tactile stimulation events that elicit a response increases significantly (binomial GLM: F_1,19_ =16.2, p<0.001, figure 2). This shows that, if a fry does respond to a tactile stimulation event, it is much more likely to initiate nudges with one or more group-mates with increasing age.

**Figure 2:**
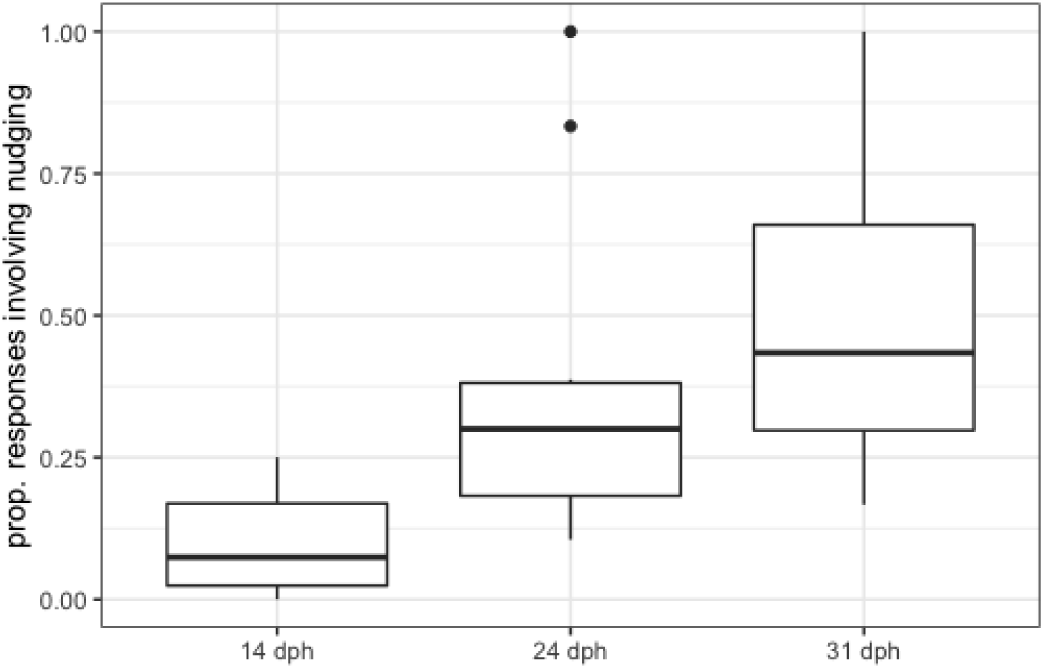
The proportion of responses involving one or more nudges (number of tactile stimulation events in which fry nudged a group-mate divided by the number of tactile stimulation events) in larvae 14, 24, and 31 days post hatching.

### The role of social exposure in responding to tactile interactions with group-mates

Groups consisting of fry raised in isolation exhibited significantly fewer nudges as compared to groups consisting of socially-reared fry (GLM: χ^2^_1_ =12.9, p<0.001, figure 3). However, there was not a significant difference in the time groups spent together based on social vs isolated housing (LM, F_1,12_= 0.88, p=0.367)

**Figure 3:**
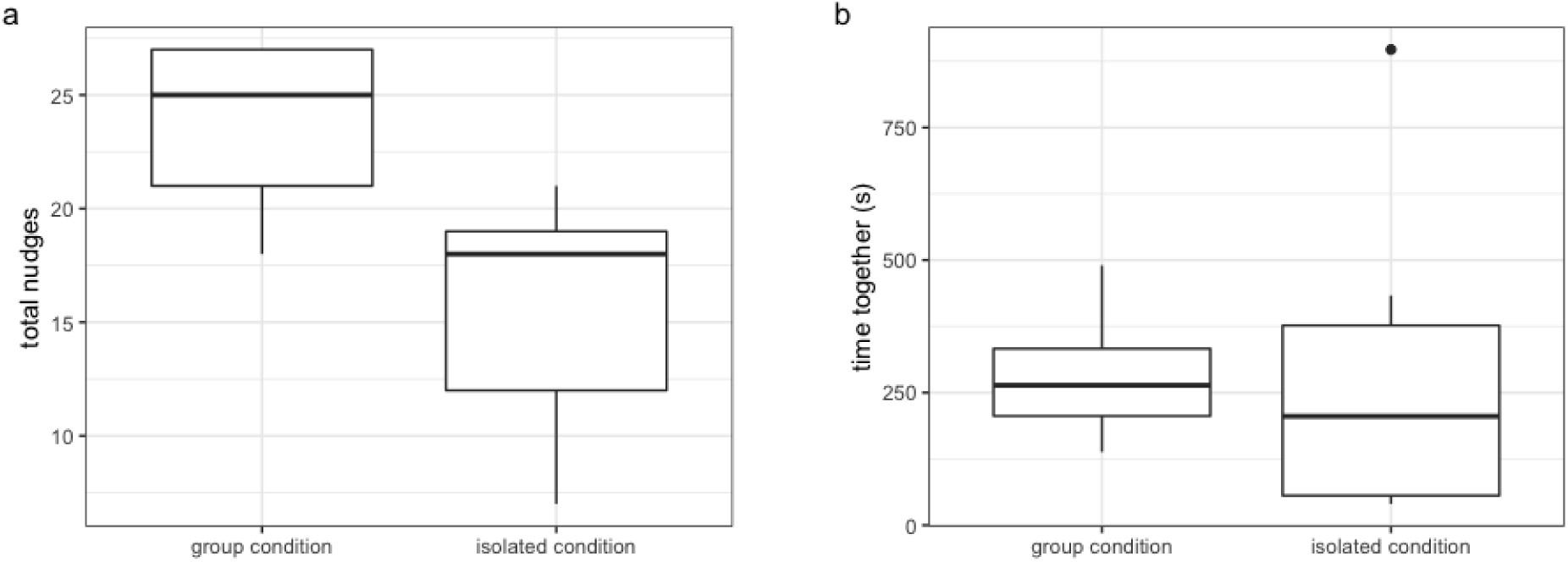
a) total nudges in groups of three fry reared socially and fry reared in isolation; b) time together in fry reared ocial and fry reared in isolation

**Figure 4:**
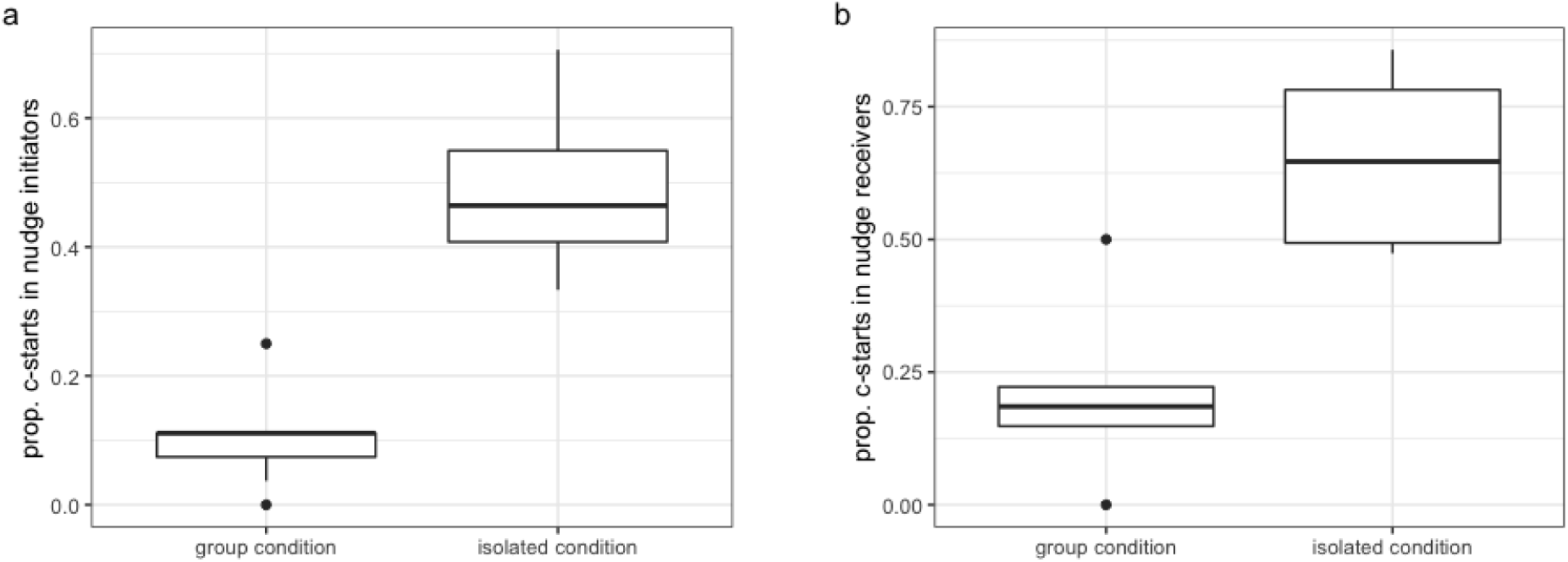
a) proportion of total interactions in which the initiator c-started following a nudge; b) proportion of total interactions in which the receiver c-started following a nudge

Fry raised in isolation are also significantly more like to respond to a nudge with a c-start both when initiating (quasibinomial GLM: F_1,12_=34.0, p<0.001) and receiving (quasibinomial GLM: F_1,12_= 21.3, p<0.001) a nudge.

However, it does not seem that individuals in the isolation condition were merely more reactive generally: the social- and isolation-reared individuals performed a similar number of spontaneous c-starts (Poisson GLM: χ^2^_1_ ==0.95, p= 0.329, figure 5).

**Figure 5:**
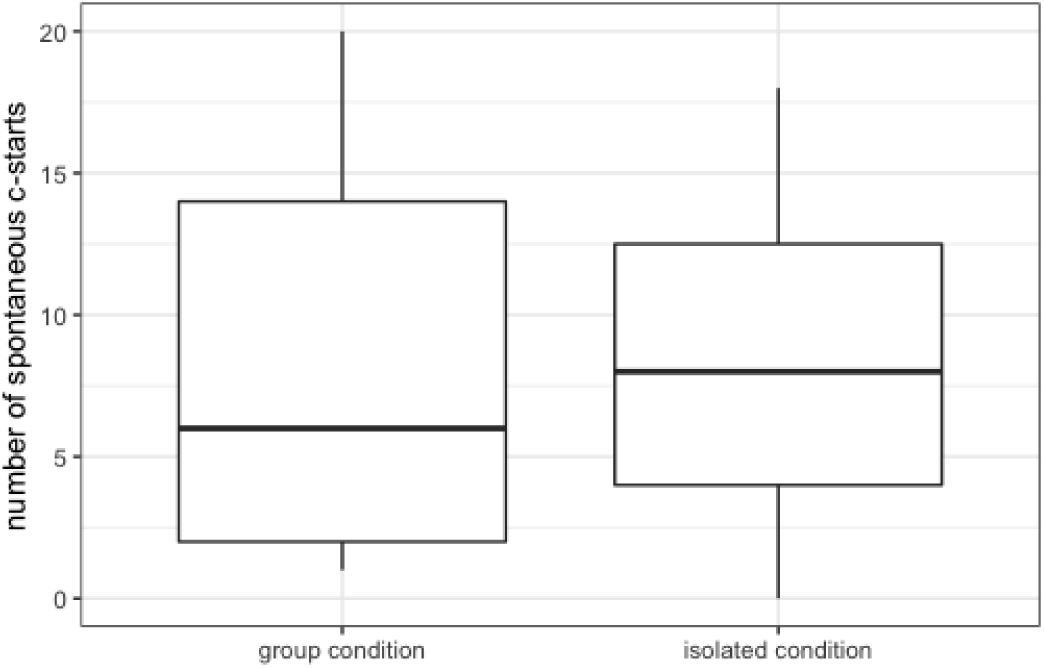
the number of spontaneous c-starts in socially- and isolation-reared groups

## Discussion

Our results show that Bronze Cory catfish larvae are more likely to c-start in response to tactile stimulation when in early developmental stages, and increasingly tolerate tactile stimulation as they develop. Our results also suggest that more developed larvae tend to initiate tactile interactions with conspecifics when disturbed by a tactile stimulation event. Increased toleration of tactile stimulation imply that the larvae may have perceived the stimulation as a threat less often with age, may have already developed the ‘freeze’ threat response common in adult threat responses, or their morphological development could have prevented them forming the c-shape necessary for a c-start to be scored.

Morphological constraints seem unlikely. Previous work on *Corydoras plateaus* found that *Corydoras* larvae are in the late pterygiolarval phase at 33 days post fertilisation (dpf) which is roughly 29dph allowing for a four-day incubation (Rodriguez-Ithurralde, 2014). Their cranial bones are still cartilaginous at 33dpf and their morphology is not very robust. It is reasonable to assume that the 31dph *C. aeneus* larvae in this study were at the same stage in development because *Corydoras plateaus* has a very similar adult size to *C. aeneus* and the larvae are a similar size for a given age (*Corydoras plateaus* 10.7mm Standard length (SL) 33dpf, *C. aeneus* 10.0mm SL 31dp). In *Corydoras arcuatus,* another similar sized species, the cartilaginous precursors to scutes do not develop until larvae are at least 15mm SL (Jean-Yves, 2005). The literature therefore suggests that morphological development at 31dph does not restrict the lateral flexion of larvae. It also seems unlikely that larvae would adopt a freeze threat response before the development of prominent anterior pectoral fin rays or armored scutes because they have very little protection from predation (Rodriguez-Ithurralde, 2014) and would rely on camouflage alone. Therefore, it seems that the most likely explanation for the change in response to tactile stimulation is that older larvae were less likely to perceive tactile stimulation as a threat. The ability to control and modulate c-start responses has been observed in archer fish in the context of feeding (Wöhl and Schuster, 2007) and it may be that the modulation of the c-start reflex occurs in Bronze Cory catfish so that tactile interaction is tolerated.

The fact that older larvae tend to initiate more tactile interactions with group-mates when they do respond to tactile stimuli is likely to be due, at least in part, to having greater social exposure with more advanced development. This is consistent with the results of the isolation experiment, in which larvae housed in groups (and which therefore have ample opportunities to interact socially with group-mates) were far less likely to respond to a tactile interaction with a group-mate with a c-start response. Social exposure has been found to influence and ultimately weaken responses to potential threats in paradise fish larvae (Ádám, 1998), as we saw with the developing larvae in our tactile stimulation experiment. It seems likely that paradise fish larvae became habituated to the continuous presence of larval conspecifics and generalized the experience of conspecifics with exposure to potential predators, leading to a weakened response to predators with increasing social exposure (Ádám, 1998). We observed a similar decrease in response to a potential threat with age in Bronze Cory catfish larvae, which was also likely a result of increased social exposure. It may be that desensitization to tactile stimulation during early development in Bronze Cories may only occur in the presence of conspecifics to ensure that individuals do not erroneously ignore a potential threat. Nonetheless, the fact that older Bronze Cory catfish larvae were more actively social (i.e. initiated more nudges with group-mates) in response to tactile stimulation implies that the effect of social exposure in Bronze Cory catfish is not just limited to the downregulation of a threat response, as it appears to be in paradise fish larvae.

Although our experiment cannot elucidate the long-term effects of social isolation, the isolated larvae’s increased likelihood of misinterpreting an interaction with a conspecific has parallels in other systems. The behavioral and neurological effects of social isolation early in development are well documented in rodent models (Fone and Porkess, 2008, Einon and Morgan, 2004, Makinodan, 2012). Impaired sensorimotor gaiting was a symptom of early social isolation in rats (Fone and Porkess, 2008) and this is one possible explanation for the higher c-start frequency we observed in isolated larvae. In many animals, including rodents (Fone and Porkess, 2008) social isolation early in development permanently alters an individual’s social behavior, rendering individuals incapable of normal social behavior or cognition. In mice, social isolation in early development has a lasting effect on brain development, but only if mice are isolated during their critical period in development (Makinodan, 2012). Because critical periods are so fundamental to the development of social behavior in many group-living animals, the study of social isolation and its effects in more diverse taxa may have the potential to improve our understanding of some psychological and behavioral disorders, and the role social exposure plays in their aetiology. Indeed, in humans, social isolation is as significant a risk factor as smoking or obesity for morbidity and mortality, effects that have also been observed in animal models (Cacioppo, 2011).

The role of social exposure in the development of adult-like behavior in Bronze Cory catfish larvae implies that there may be a developmentally vital period during which individual social behavior develops through social exposure. In the tactile stimulation experiment, the decrease in c-start frequency seemed to occur gradually over time, with 14 dph larvae c-starting frequently, 24 dph larvae c-starting significantly less frequently, and 31 dph larvae c-starting very seldomly at all. This progression seems gradual over this time period, without a developmental ‘switch’ that very rapidly modifies behavior, but this window of time may represent a critical period for social skill acquisition. Although this gradual decrease could be the result of acclimation to the tactile stimulus, the result that isolated larvae in the isolation experiment exhibited significantly more c-starts in response to nudges from conspecifics strongly suggests that sustained social exposure leads to the development of toleration for tactile stimulation.

Although our isolation experiment provides evidence that nudging requires social exposure to properly develop, it is unclear whether or not there is a critical period of social exposure for the development of nudging. In our isolation experiment, the isolation of larvae was confounded when they formed groups of three for filming and the development of isolated individuals was not followed post filming due to the difficulty of tagging larvae. Future work may aim to determine if the effect of social isolation we observed in isolated larvae can be overcome by social exposure later in life or if Bronze Cory catfish larvae show a critical period for social exposure. Social behavior may be plastic throughout a fish’s life or it could be determined by early developmental environment. If sociality in the Bronze Cory catfish does display a critical period for development, then it is ecologically very important for an individual to have social exposure and interact with other larvae during that critical period. If a larva does not, then it may become unable to interact effectively with the rest of the shoal later in life, potentially misinterpreting the social interactions of conspecifics as a threat. Their readily observable tactile interaction behavior and response paradigm make the Bronze Cory catfish a compelling model system for investigating the development of social behavior and its consequences in later life.

## References

Ádám, M., Péter, P., Vilmos, C., 1998. The ontogeny of antipredator behavior in paradise fish larvae (Macropodus opercularis) IV. The effect of exposure to siblings. Dev. Psychobiol. 30, 283–291. https://doi.org/10.1002/(SICI)1098-2302(199705)30:4<283::AID-DEV2>3.0.CO;2-K

Buske, C., Gerlai, R., 2011. Shoaling develops with age in Zebrafish (Danio rerio). Prog. Neuropsychopharmacol. Biol. Psychiatry 35, 1409–1415. https://doi.org/10.1016/j.pnpbp.2010.09.003

Cacioppo, J.T., Hawkley, L.C., Norman, G.J., Berntson, G.G., 2011. Social isolation. Ann. N. Y. Acad. Sci. 1231, 17–22. https://doi.org/10.1111/j.1749-6632.2011.06028.x

Eaton, R.C., DiDomenico, R., Nissanov, J., 1988. Flexible body dynamics of the goldfish C-start: implications for reticulospinal command mechanisms. J. Neurosci. 8, 2758 LP – 2768.

Eaton, R.C., Emberley, D.S., 1991. How stimulus direction determines the trajectory of the Mauthner-initiated escape response in a teleost fish. J. Exp. Biol. 161, 469 LP – 487.

Einon, D.F., Morgan, M.J., 2004. A critical period for social isolation in the rat. Dev. Psychobiol. 10, 123–132. https://doi.org/10.1002/dev.420100205

Fone, K.C.F., Porkess, M.V., 2008. Behavioural and neurochemical effects of post-weaning social isolation in rodents—Relevance to developmental neuropsychiatric disorders. Neurosci. Biobehav. Rev. 32, 1087–1102. https://doi.org/10.1016/j.neubiorev.2008.03.003

Jean-Yves, S., 2005. Development and fine structure of the bony scutes in Corydoras arcuatus (Siluriformes, callichthyidae). J. Morphol. 215, 225–244. https://doi.org/10.1002/jmor.1052150305

Kimmel, C.B., Eaton, R.C., Powell, S.L., 1980. Decreased fast-start performance of zebrafish larvae lacking mauthner neurons. J. Comp. Physiol. 140, 343–350. https://doi.org/10.1007/BF00606274

Kohda, M., Tanimura, M., Kikue-Nakamura, M., Yamagishi, S., 1995. Sperm drinking by female catfishes: a novel mode of insemination. Environ. Biol. Fishes 42, 1–6. https://doi.org/10.1007/BF00002344

Magurran, A.E., 1990. The adaptive significance of schooling as an anti-predator defence in fish. Ann. Zool. Fenn. 27, 51–66.

Makinodan, M., Rosen, K.M., Ito, S., Corfas, G., 2012a. A Critical Period for Social Experience–Dependent Oligodendrocyte Maturation and Myelination. Science 337, 1357 LP – 1360.

Makinodan, M., Rosen, K.M., Ito, S., Corfas, G., 2012b. A Critical Period for Social Experience–Dependent Oligodendrocyte Maturation and Myelination. Science 337, 1357–1360. https://doi.org/10.1126/science.1220845

Pitcher, T.J., Magurran, A.E., Winfield, I.J., 1982. Fish in larger shoals find food faster. Behav. Ecol. Sociobiol. 10, 149–151. https://doi.org/10.1007/BF00300175

Riley, Riva J., Gillie, E.R., Johnstone, R.A., Boogert, N.J., Manica, A., 2018. Coping with strangers: how familiarity and active interactions shape group coordination in Corydoras aeneus. bioRxiv 448068. https://doi.org/10.1101/448068

Riley, Riva J, Gillie, B.R., Jungwirth, A., Savage, J., Boogert, N.J., Manica, A., 2018. The role of tactile interactions in flight responses in the Bronze Cory catfish (Corydoras aeneus). bioRxiv. https://doi.org/10.1101/449272

Rodríguez-Ithurralde, D., del Puerto, G., Fernández-Bornia, F., 2014. Morphological development of Corydoras aff. paleatus (Siluriformes, Callichthyidae) and correlation with the emergence of motor and social behaviors. Iheringia. Série Zoologia 104, 189–199. https://doi.org/10.1590/1678-476620141042189199

Slagsvold, T., Hansen, B.T., Johannessen, L.E., Lifjeld, J.T., 2002. Mate choice and imprinting in birds studied by cross-fostering in the wild. Proc. R. Soc. Lond. B Biol. Sci. 269, 1449–1455. https://doi.org/10.1098/rspb.2002.2045

Spence, R., Smith, C., n.d. The Role of Early Learning in Determining Shoaling Preferences Based on Visual Cues in the Zebrafish, Danio rerio. Ethology 113, 62–67. https://doi.org/10.1111/j.1439-0310.2006.01295.x

Weihs, D., 1973. The mechanism of rapid starting of slender fish. Biorheology 10, 343–350.

Wöhl, S., Schuster, S., 2007. The predictive start of hunting archer fish: a flexible and precise motor pattern performed with the kinematics of an escape C-start. J. Exp. Biol. 210, 311 LP – 324.

